# Electric shock causes a fear-like persistent behavioral response in the nematode *Caenorhabditis elegans*

**DOI:** 10.1101/2021.09.07.459218

**Authors:** Ling Fei Tee, Jared J. Young, Ryoga Suzuki, Keisuke Maruyama, Yuto Endo, Koutarou D. Kimura

## Abstract

Behavioral persistency reflects internal brain states, which are the foundations of multiple brain functions. However, experimental paradigms that enable genetic analyses of behavioral persistency and its associated brain functions have been limited. Here we report novel persistent behavioral responses caused by electric stimuli in the nematode *Caenorhabditis elegans*. When the animals on bacterial food are stimulated by alternating current, their movement speed suddenly increases more than 2-fold, which persists for minutes even after the electric stimulation is terminated. Genetic analyses reveal that multiple types of voltage-gated channels are required for the response, possibly as the sensors, and neuropeptide signaling regulates the duration of the persistent response. Additional behavioral analyses indicate that the animal’s response to electric shock is scalable and has a negative valence. These properties, along with persistence, have been recently regarded as essential features of emotion, suggesting that the animal’s response to electric shock may express a form of emotion, such as fear.

## INTRODUCTION

Animal behaviors, such as feeding, mating, aggression, and sleeping, are strongly related to internal states in the brain, namely motivation, arousal, drive, and emotion (Anderson, 2016; Berridge, 2004; Kennedy et al., 2014). Because animals can produce different behavioral responses to the same stimulus depending on their brain state, these states are considered to be the foundation from which a variety of behavioral responses emerge (Chen & Hong, 2018; Maimon, 2011). The brain states persist for a certain period of time and transit to a different state based on internal and/or external triggers, which can be observed as transitions among different persistent behavioral states. The neural mechanisms of brain/behavioral states are starting to be revealed. For example, the behavioral states of mating and aggressiveness have been shown to be controlled by relatively small circuits in mice and flies (Anderson, 2016; Hoopfer et al., 2015; Lee & Dan, 2012). However, the mechanisms of persistent brain/behavioral states have been revealed in only limited studies, and, moreover, the molecular basis that generates persistent states is still unclear.

The nematode *Caenorhabditis elegans* has been widely used in neurobiological research because of the feasibility of molecular, physiological, and behavioral analyses of neural functions (Bargmann, 2006; de Bono & Maricq, 2005; Sasakura & Mori, 2013). Recently, persistent behavioral states have also been studied in the animals, especially roaming/dwelling and sleep/arousal. Roaming/dwelling are states of locomotion on bacterial food, either moving over long distances at a constant speed or moving back and forth over short distances (Ben Arous et al., 2009; Fujiwara et al., 2002). Sleep in *C. elegans* is a phenomenon observed just before molt, and meets the definition of sleep in higher animals such as humans, rodents, fishes and flies (Raizen et al., 2008). Both the neural circuits and genes that control these phenomena are being revealed (Flavell et al., 2020). However, it is not clear whether *C. elegans* has any other persistent states.

In this study, we report that *C. elegans* exhibits a novel type of persistent behavioral response to electric stimulus. The animals respond to alternating current (AC) stimulus by immediately increasing their speed, and the speed increase persists even for minutes after five seconds of electric stimulus: This result suggests that the response is caused not by direct stimulation of the motor system for rapid movement but by persistent activity of a specific set of neurons to generate the behavioral response. Further behavioral analyses indicate that the speed increase to AC stimulus is scalable and has negative valence. Because persistent behavioral response is one of the most prominent characteristics of emotions of animals (Abbott, 2020; Anderson & Adolphs, 2014; Nettle & Bateson, 2012; Paul & Mendl, 2018; Perry & Baciadonna, 2017), and persistency, scalability and valence are three of the four key features of animal emotions proposed by Anderson and Adolphs (Anderson & Adolphs, 2014), the speed increase to the electric shock may express a form of emotion. A series of candidate genetic analyses reveal that the response is not mediated by well-known chemo- or mechano-sensory mechanisms. Instead, it requires voltage-gated calcium and potassium channel genes, which are required for electro-sensation in cartilaginous fishes (Bellono et al., 2017, 2018), suggesting an evolutionarily conserved mechanism for electro-sensation. Furthermore, we find that neuropeptide signaling regulates the duration of persistence. These results indicate that the animals’ response to electric shock can be a suitable paradigm to reveal molecular and physiological mechanisms of electro-sensation as well as persistent brain/behavioral states.

## RESULTS

### *C. elegans*’ speed is increased by AC stimulation

Initially, we started this project by studying *C. elegans* behavioral responses to AC stimuli. The animals are known to respond to direct current (DC), migrating along the electric field from the positive end to the negative end (Sukul & Croll, 1978), and a few classes of chemosensory neurons (ASH and ASJ) were found to be required for their ability to align themselves according to the DC field (Gabel et al., 2007). However, the animal’s migratory response to AC stimulus has not been reported yet. In our original setup (Fig. 1), several adult wild-type animals were placed onto 9 cm agar plates seeded with a small bacterial food patch and subjected to AC stimulation. The complete trajectories produced by the animals were video-recorded, and their speed was calculated.

**Fig. 1.**
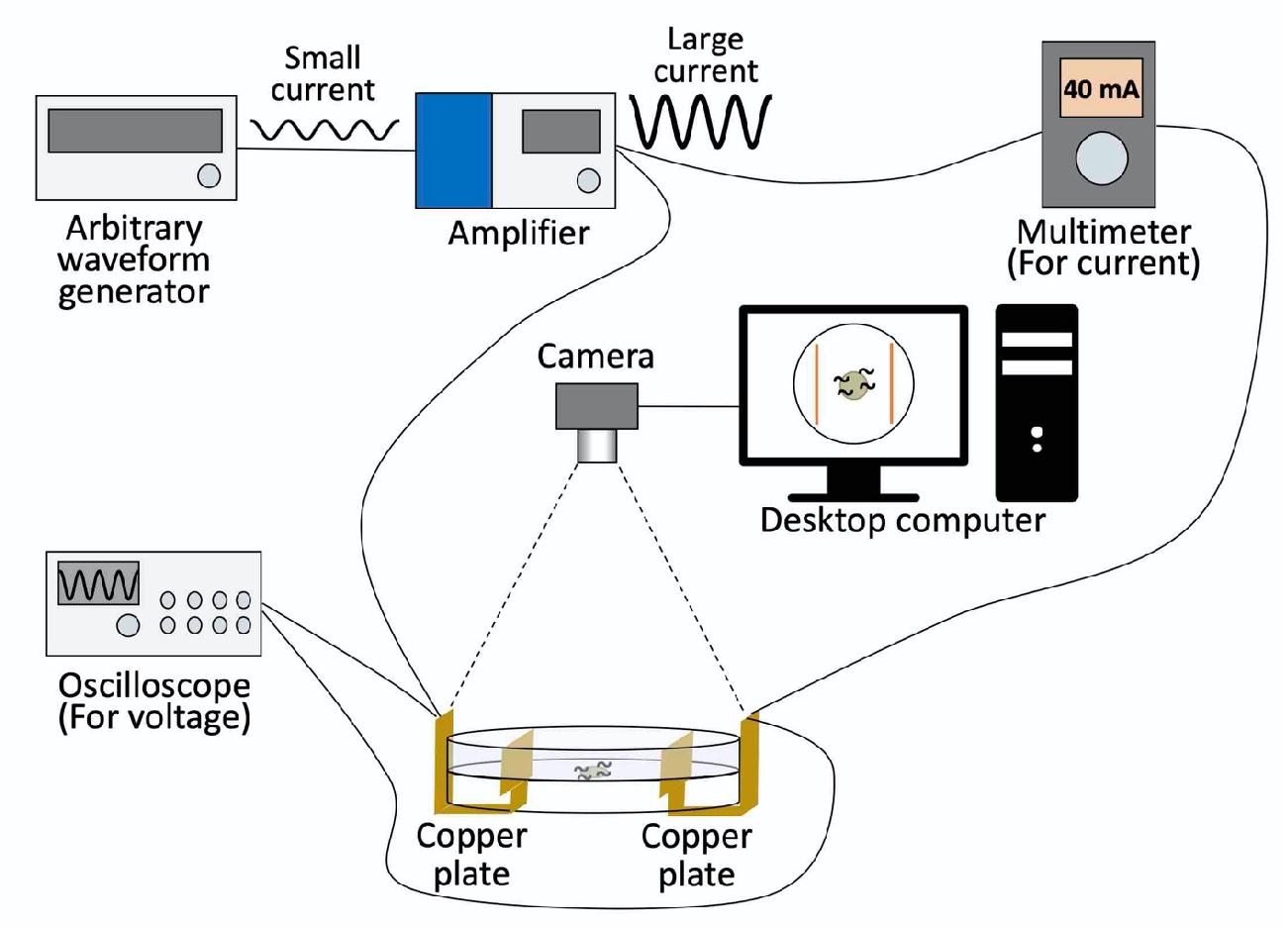
Experimental setup of electric shock experiment. This setup consists of an arbitrary waveform generator, amplifier, multimeter, oscilloscope, camera, and desktop computer.

We first studied the response to AC stimulation covering a range between 15 - 105 V at 60 Hz (the commercial power frequency in Japan), and found that the animals increased their average speed during electric stimulation by varying amounts (Supplementary Fig. 1). We then conducted a series of systematic analyses with different voltages and frequencies at 30– 75 V and 0.25–256 Hz (Supplementary Fig. 2). After the analysis, we noticed that an interesting characteristic of this behavioral phenotype is most apparent when using 4 Hz stimuli: When animals were stimulated with 30 V, their average speed of movement suddenly increased more than 2-fold, and this persisted during the electric admission. We named this behavior the “ON response” (Fig. 2A and C). During this running behavior, the animals engage in rapid body bends as well as rapid head movements (Supplementary Videos 1 and 2). In the ON response, we did not detect a statistical bias in direction (Supplementary Fig. 3). Unexpectedly, when a stronger electric stimulus of 75 V was applied, it caused a significant increase in average speed not during but immediately after the stimulus, which we named the “OFF response” (Fig. 2B). A fraction of the animals responded during the stimulus in the OFF response condition, while, in the majority of the animals, the speed was suppressed during the stimulus and then increased immediately after its removal (Supplementary Fig. 4 and Videos 3 and 4). With other frequencies, ON and OFF responses were also observed, but were less clear compared to those with 4 Hz (Supplementary Fig. 2). The range of voltage per length (30–75 V/6 cm = 5–12.5 V/cm) is similar to the range previously shown to elicit responses to DC (3–12 V/cm) (Gabel et al., 2007), suggesting that these electric stimuli are physiologically meaningful for the animals. The fact that ON and OFF responses at 4 Hz were substantially different with only a 2.5-fold difference in the voltage at the same frequency is interesting because different behavioral responses generally require much larger differences in stimulus intensity with other stimuli, such as odor (Bargmann et al., 1993).

**Fig. 2.**
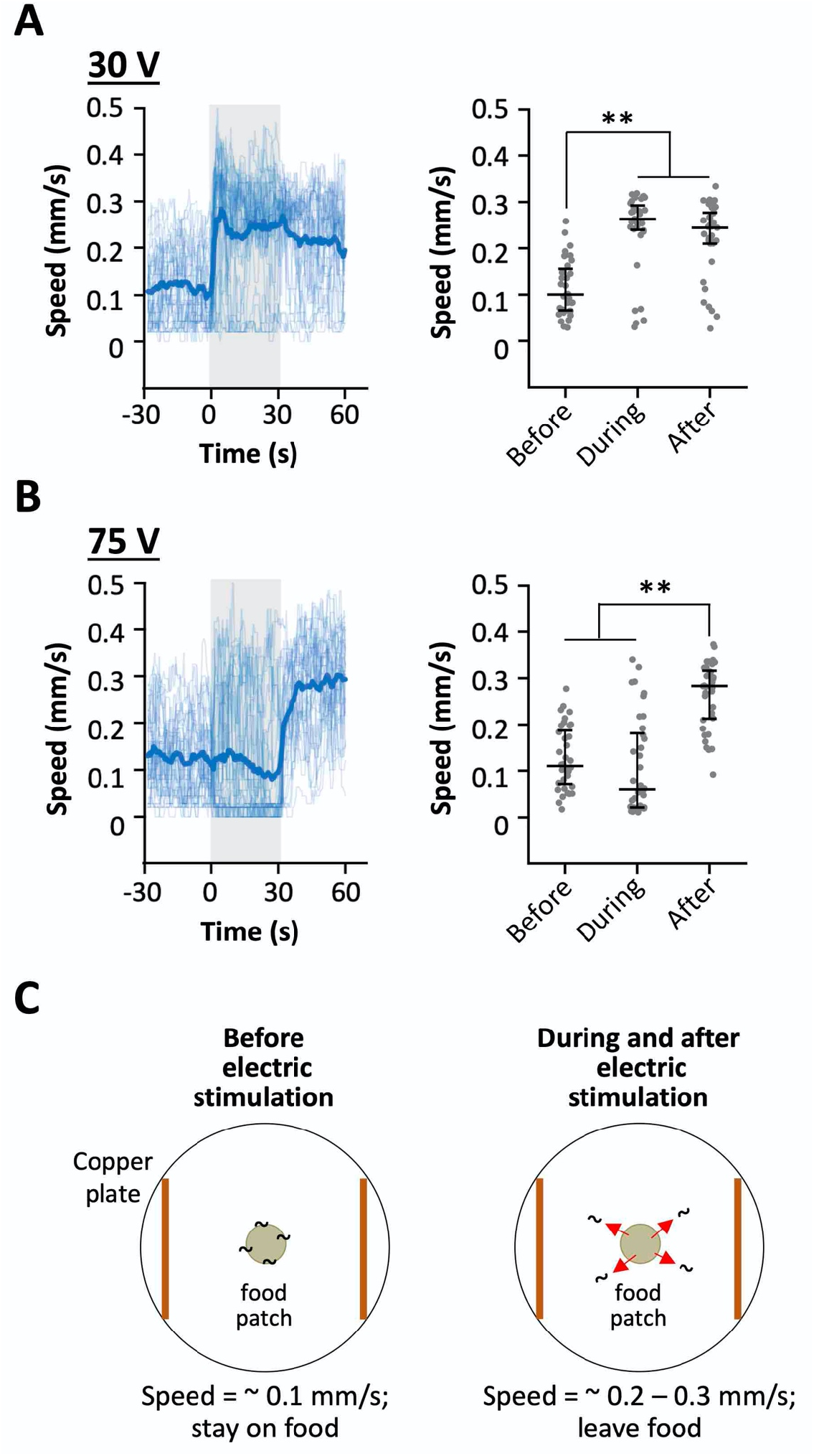
Animals’ speed is increased by AC stimulation. **A,** (Left) Speed-time graph with 30 V stimulation at 4 Hz. Thin and thick lines are for individual and average values, respectively. Gray indicates the duration of electric stimulation (0-30 s). (Right) Scatter plot showing average speed of individual animals before, during and after electric stimulation. Each period is 30 s. n = 35. **B,** Speed-time graph (left) and scatter plot (right) with 75 V stimulation at 4 Hz. n = 36.
**C,** Cartoons of worm’s response to the electric shock. (Left) Before electric stimulation, the animals stay on food patch and maintain their speed at around 0.1 mm/s. (Right) During electric stimulation is delivered, the animals increase speed to around 0.2 - 0.3 mm/s and leave the food patch which persists even after the stimulus is terminated. Statistical values were calculated using Kruskal-Wallis test with Bonferroni correction. ** *p* < 0.001.

We then analyzed whether this response depends on voltage or current by manipulating the salt concentration in the assay plate (Fig. 3): When 30V is applied to the high salt plate the current should be similar to the current produced when 75V is applied to a plate with our standard (control) salt concentration. Conversely, when 75V is applied to the low salt plate, the current should be similar to the current produced when 30V is applied to the control plate (Fig. 3A). As shown in Fig. 3, 30 V and 75 V stimuli caused ON and OFF responses, respectively, regardless of the current value, indicating that the behavioral response depends on voltage.

**Fig. 3.**
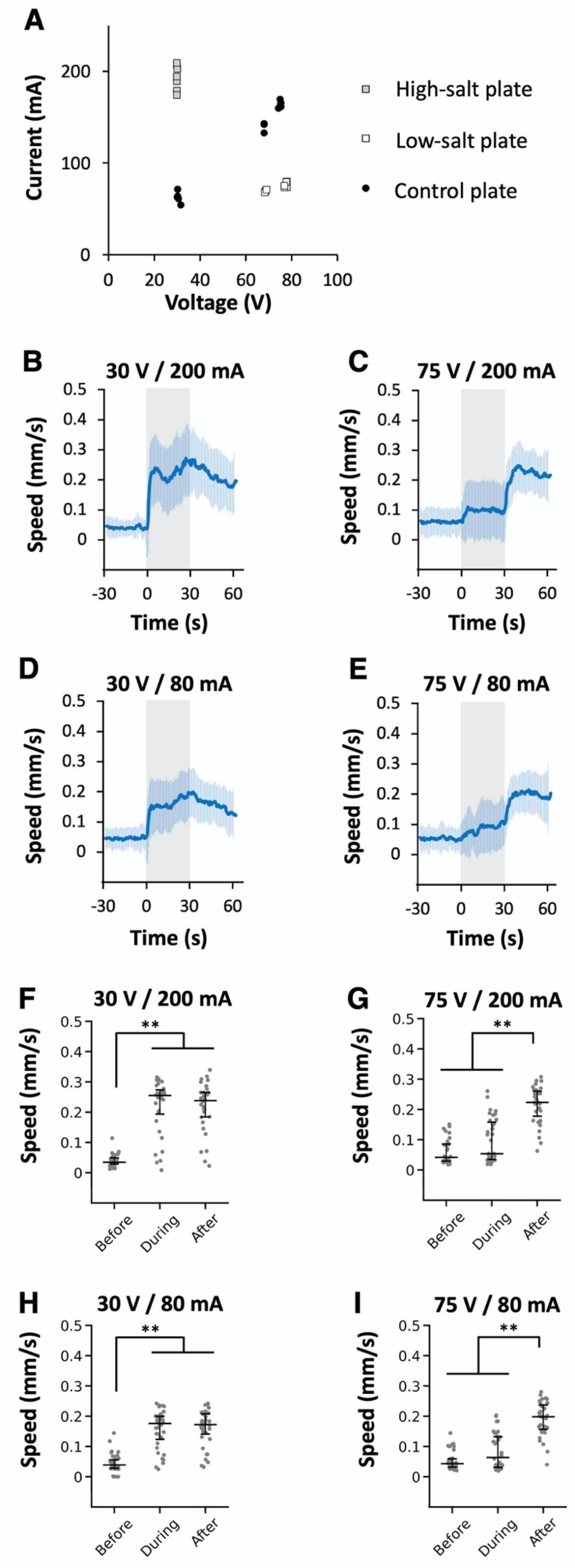
Speed increase is dependent on voltage, not on current. **A,** Voltage-current graph with different salt concentrations (indicated by different symbols). Each dot represents the measured value on the day of the experiment. The final total osmolarity for sodium chloride (Na^+^ and Cl^-^) and sucrose for all the plates was 400 mOsm. **B–E,** Behavioral responses of animals assayed on high-salt plate with 30 V (**B**; n = 32), on control plate with 75 V (**C**; n = 35) or 30 V (**D**; n = 36), or on low-salt plate with 75 V (**E**; n = 34). Stimulation period is indicated by a shaded grey box. **F–I,** Scatter plot showing average speed of individual animals before, during and after electric stimulation, corresponding to the panels **B–E**, respectively. Statistical values were calculated using Kruskal-Wallis test with Bonferroni correction. ** *p* < 0.001.

### Speed increase lasts for several minutes

Next, we examined how long the increased speed persists during and after the stimulus. When the duration of 30V applied electric shock was 1-2 minutes, significant speed increases were maintained during the stimulus, lasted for ~1 min after the stimulus, and went back to the baseline level (Fig. 4A). Interestingly, when the animals were stimulated only for 5 sec, the speed increase still lasted for 1.5 min. When 4 min stimulus was applied, the increase was maintained during the stimulus but went back to the baseline level 30 sec after the stimulus. During 10 min stimulation, the significant speed increase was observed only for 5.5 min. Thus, we concluded that the ON response caused by 30 V stimulation persists ~5 min at most.

**Fig. 4.**
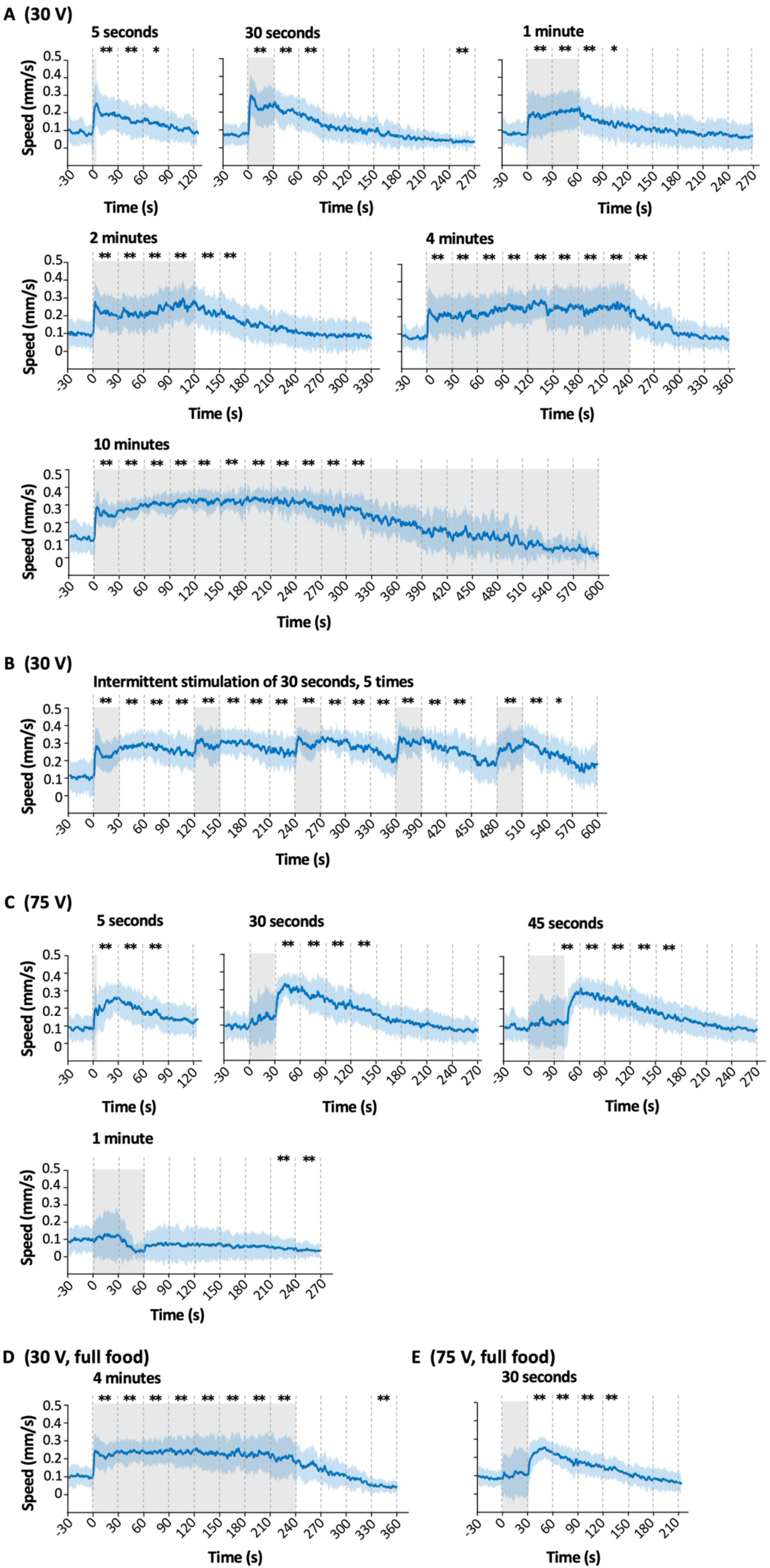
Speed increase persisted for minutes even after the stimulation. **A,** Speed-time graphs of ON response with 30 V stimulation of different time periods, ranging from 5 seconds to 10 minutes. **B,** Speed-time graph for intermittent electric stimulation of 30 seconds, 5 times with 90 s–intervals. **C,** Speed-time graphs of OFF response with 75 V stimulation of different time periods, ranging from 5 seconds to 1 minute. **D** and **E,** Speed-time graphs for electric stimulation of 30 V for 4 minutes (**D**) or 75 V for 30 s (**E**) with animals placed on full food lawn. Shaded regions around the lines represent standard deviation. Statistical values were calculated using Kruskal-Wallis test with Bonferroni correction for the differences from the average speed before the stimulation. * *p* < 0.01, ** *p* < 0.001. Sample numbers were 32–46 per condition, and the details are described in Supplementary Table.

This result suggested the possibility that the speed increase may decline after several minutes because of fatigue in motor systems. However, animals stimulated intermittently 5 times for 30 seconds per stimulation maintained a speed increase for a much longer time than those under the continuous stimulus (Fig. 4B versus “10 minutes” in A). This result supports the idea that the decrease in speed during the long ON stimulation period is not caused by fatigue in the motor system, but possibly by sensory adaptation, which is widely known to adjust the animal’s sensory response to new environments (Wark et al., 2007).

We then tested the persistence of speed increase in the OFF response with 75 V. Five and 30 sec stimuli caused similar or longer persistent responses after the stimulus than 30 V did (Fig. 4C). Remarkably, 45 sec stimulus caused >2 min persistent response, which is the longest among the responses to 30 and 75 V stimuli after the stimulus. When animals were stimulated for 1 min, no ON or OFF responses were observed. The fact that the larger stimulus (75 V) caused longer persistent responses than the smaller one (30 V) suggests that the response to electric shock is “scalable” (i.e., different strength of stimulus causes different strength of behavioral response), one of the critical “emotion primitives” together with persistence (Anderson & Adolphs, 2014).

We then tested the effect of food presence on the speed increase. *C. elegans* move slowly on the bacterial food lawn and faster out of the lawn (Sawin et al., 2000). As we used a small food lawn to localize the animal’s initial positions to the center of the plate (Fig. 1 and 2C), it was possible that the electric stimulus caused the animals to move away from the food lawn, which then caused increased speed due to the absence of food. If this is the case, the animal’s speed would be considerably lower with the electric stimulus when the plates were fully covered with a bacterial lawn. To test this hypothesis, we compared the time-course of speed changes on plates with a small patch of food lawn and with a full food lawn. As shown in Fig. 4D and E (compare Fig. 4A “4 minutes” and C “ 30 seconds”, respectively), there was no substantial difference in the time course of speed change between the small food and the full food plates in ON as well as OFF responses, demonstrating that the speed increase is not caused by the food absence but by the electric stimulation itself.

To further confirm that result, we analyzed the animals’ speed on a stripe-like food pattern (Supplementary Fig. 5A). We did not observe a significant difference in speeds when the animals moved into or out of the food area (Supplementary Fig. 5B). This result suggests that the electric stimulus may have negative valence that is more influential to the animal’s behavior than the food signal, even though food is critical for their survival. It further suggests that animals prioritize moving away from a harmful condition, such as the electric shock, to protect themselves.

### Voltage-gated ion channel genes and neuropeptide signaling are required for the AC response

The molecules required for responses to electric signals have only been revealed in cartilaginous fishes: Bellono et al. reported that electrosensory cells in little skate and chain catshark use L-type voltage-gated calcium channels (VGCC) and voltage-gated big-conductance potassium (BK) channels (Bellono et al., 2017, 2018). To identify gene(s) required for the response to electric shock in *C. elegans*, we analyzed a series of mutant strains of candidate genes. Specifically, we tested mutants of genes involved in the animals’ chemo- and mechano-sensation, and the homologues of genes involved in electroreception in the cartilaginous fishes.

*C. elegans*’ chemo-sensation is largely mediated by the 12 pairs of amphid sensory neurons in the head, which are classified into the ones using TAX-2 and TAX-4 cyclic nucleotide-gated channel (CNGC) subunits or the others using OSM-9 and OCR-2 transient receptor potential (TRP) channel subunits for depolarization (Coburn & Bargmann, 1996; Colbert et al., 1997; Komatsu et al., 1996; Tobin et al., 2002). In addition to loss-of-function mutants for the above-mentioned genes, we tested mutants for *che-2*, a gene required for the proper formation and function of the sensory cilia (Fujiwara et al., 1999). For mechano-sensation, we analyzed loss- or reduction-of-function alleles of *mec-4, mec-10*, and *trp-4. mec-4* and *mec-10* genes encode DEG/ENaC proteins that form a mechanosensory ion channel complex for transduction of gentle touch (Driscoll & Chalfie, 1991; Huang & Chalfie, 1994), while *trp-4* encodes TRPN (NOMPC) for harsh touch response (Kang et al., 2010). All these mutant strains exhibited wild-type-like ON and/or OFF responses (panel A in Fig. 5 and 6 for ON and OFF responses, respectively). Some mutants (*osm-9;ocr-2, che-2, mec-4, mec-10*, and *tph-1*) exhibited statistical differences in either the ON or OFF response, but not both (Fig. 5F and 6F), suggesting the partial involvement of these genes, although the defects in speed increase (i.e. ΔSpeed) were not as severe as the ones of VGCC mutants (see below). The non-involvement of *tax-4* also indicates that temperature increase caused by the electric stimulus is not responsible for the speed increase (see Discussion for details).

**Fig. 5.**
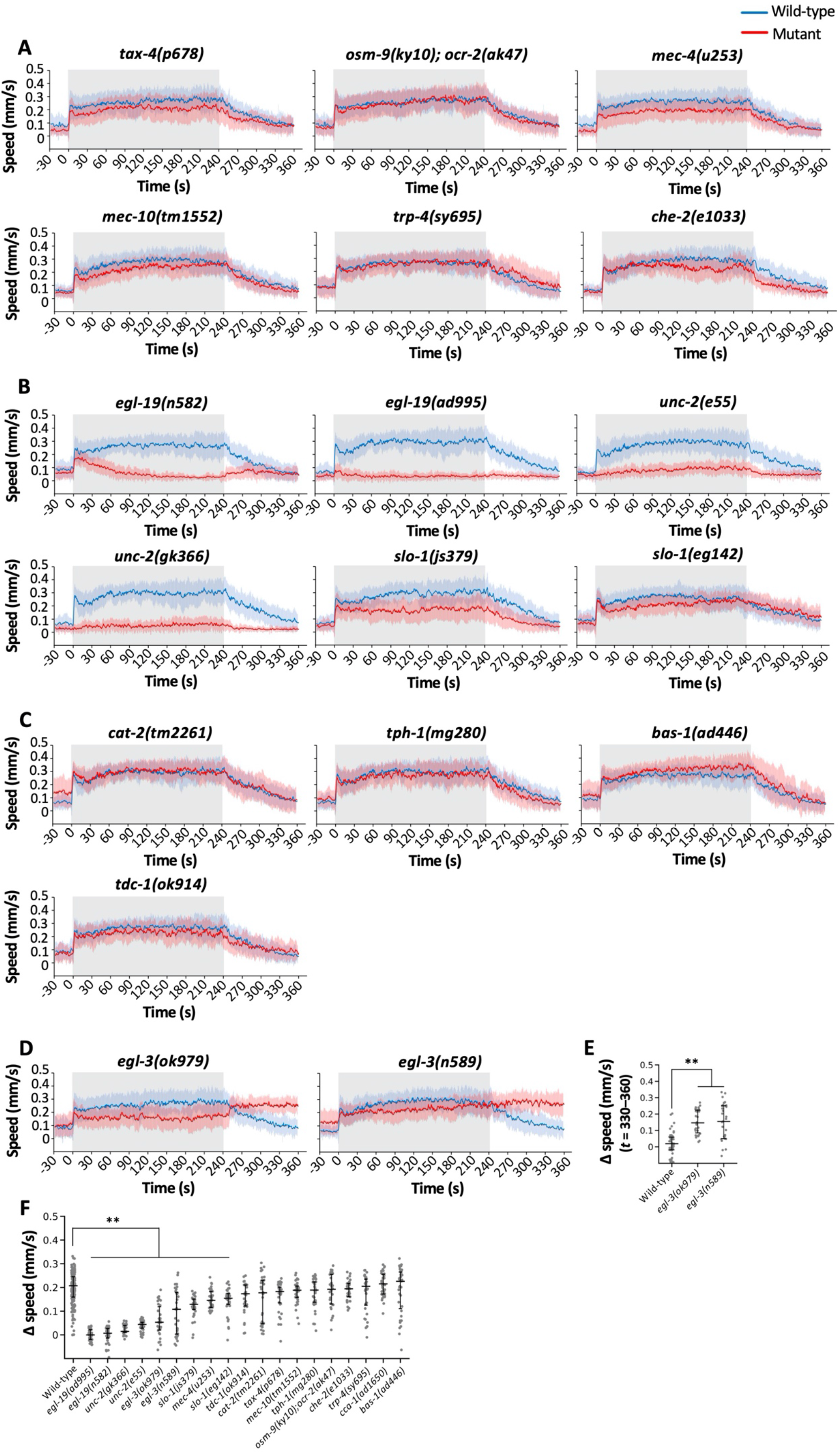
Genetic analysis of ON response. **A–D,** Speed-time graphs of ON response with 30 V stimulation of 4 min on mutants of sensory signaling (**A**), voltage-gated channels (**B**), biogenic amine biosynthesis (**C**), and neuropeptide biosynthesis (**D**). **E**, Scatter plot showing Δspeed of individual wild-type and *egl-3* mutant animals during *t* = 330-360 s in **D**. **F**, In a series of daily experiments, wild-type N2 and three to five mutant strains were analyzed in parallel. All the N2 data are combined, and the mutant strains are arranged in ascending order of median values in **F**. Statistical values were calculated using Kruskal-Wallis test with Bonferroni correction. ** *p* < 0.001. Sample numbers were 30–36 per mutant strain, and the details are described in the Supplementary Table.

**Fig. 6.**
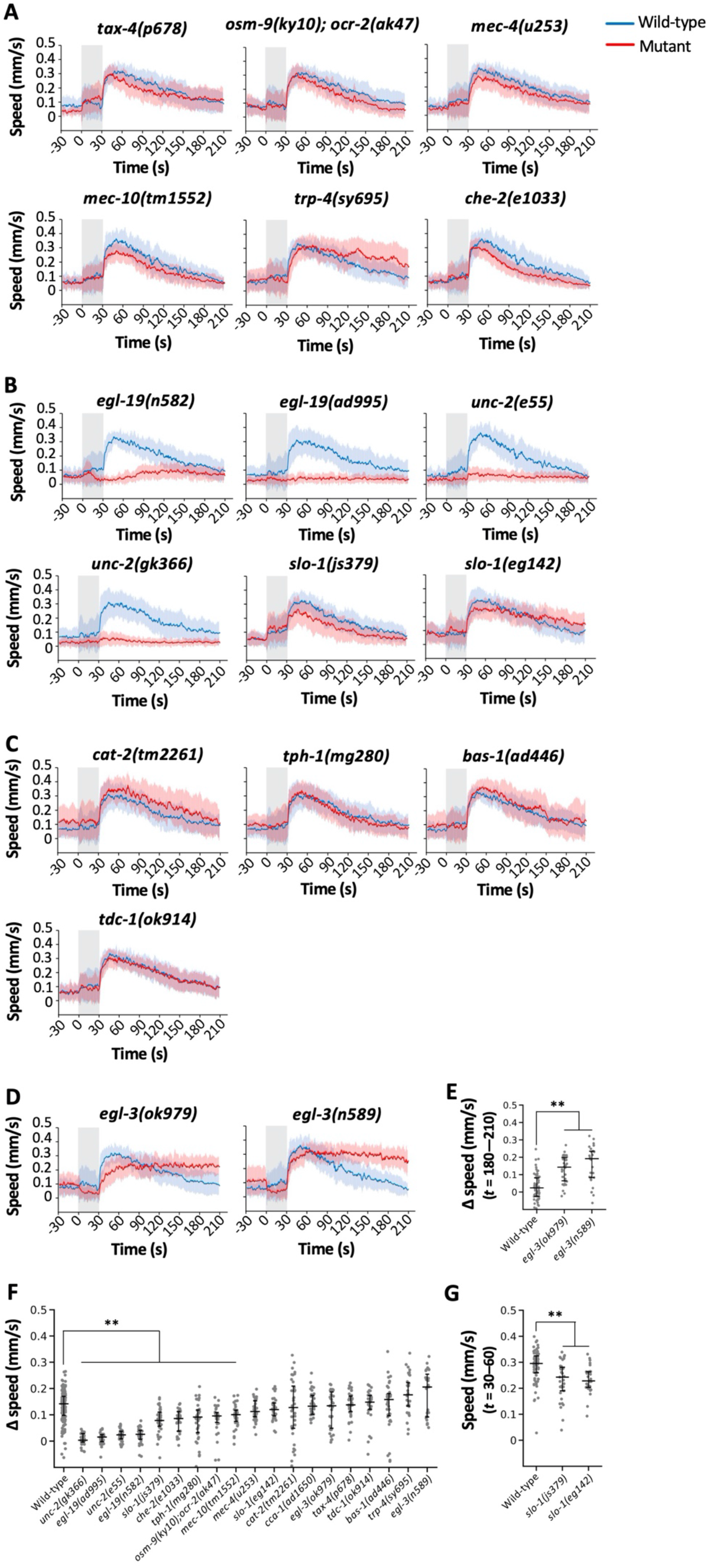
Genetic analysis of OFF response. **A–D,** Speed-time graph of OFF response with 75 V stimulation of 30 s on mutants of sensory signaling (**A**), voltage-dependent channels (**B**), biogenic amine biosynthesis (**C**), and neuropeptide biosynthesis (**D**). **E**, Scatter plot showing Δspeed of individual wild-type and *egl-3* mutant animals during *t* = 180-210 s in **D**. **F**, Scatter plot showing Δspeed of individual animals during 3 min after the stimulation (*i.e*., *t* = 30-210 s). In a set of daily experiments, wild-type and three to five mutant strains were analyzed in parallel. All the wild-type data are combined, and the mutant strains are arranged in ascending order of median values in **F**. **G**, Scatter plot showing average speed of individual wild-type and *slo-1* mutant animals during 30 s after the stimulation (*i.e., t* = 30-60 s). Statistical values were calculated using Kruskal-Wallis test with Bonferroni correction. ** *p* < 0.001. Sample numbers were 30–36 per mutant strain, and the details are described in the Supplementary Table.

We then tested *egl-19*, the orthologue of the L-type VGCC alpha subunit (Lee et al., 1997), which functions in the sensory organ for environmental electric signals for cartilaginous fishes (Bellono et al., 2017, 2018). We found that two reduction-of-function alleles of *egl-19* mutants exhibited strong defects in ON and OFF responses (Fig. 5 and 6, panels B and F). While *egl-19* is expressed widely (Lee et al., 1997), at least one allele of *egl-19* mutants exhibited movement speed comparable to wild-type animals before stimulation (Supplementary Fig. 6), suggesting that the defect in the response is not caused by a problem in the basal locomotory system. These results suggest that the VGCC may be an evolutionarily conserved sensor for environmental electricity.

This finding further motivated us to test two other types of voltage-gated calcium channels, namely, N-type (UNC-2) and T-type (CCA-1) VGCCs (Schafer & Kenyon, 1995; Steger et al., 2005), although only L-type VGCC had been found to be involved in electrical responses in the cartilaginous fishes. Unexpectedly, mutants for two alleles of *unc-2* were defective in both ON and OFF responses, while *cca-1* mutants behaved similar to the wild-type controls (Fig. 5 and 6, panels B and F, and Supplementary Fig. 7).

We then investigated the involvement of the BK channel, a voltage-gated potassium channel, also known to be involved in electro-sensation in cartilaginous fish (Bellono et al., 2017, 2018). Interestingly, two alleles of *slo-1*, the sole ortholog of BK channels in *C. elegans* (Davies et al., 2003; Wang et al., 2001), also exhibited statistical differences in the ON as well as the OFF response, at least in some aspects (Fig. 5B and F for ON response, and 6B, F and G for OFF response). The possible involvement of BK channels in addition to the VGCC in the *C. elegans*’ electrical response suggest that the molecular mechanisms of electro-sensation may be evolutionally conserved.

Lastly, we attempted to identify genes required for the behavioral persistency, and considered the genes involved in the biosynthesis of neuromodulators. We tested *cat-2* (dopamine), *tph-1* (serotonin), *bas-1* (dopamine and serotonin) and *tdc-1* (tyramine and octopamine) mutant animals (Alkema et al., 2005; Lints & Emmons, 1999; Loer & Kenyon, 1993; Sze et al., 2000), and all of these mutants exhibited wild-type-like responses, indicating that these neuromodulators are not involved (panel C in Fig. 5 and 6). Because dopamine and serotonin signaling are known to be required for the feeding status-dependent modulation of migratory speed, these results are also consistent with the fact that feeding status is not the causal reason for the speed increase (Fig. 4D and E, and Supplementary Fig. 5).

We then further tested the involvement of neuropeptides by using loss- or reduction-of-function mutations of *egl-3*, a gene required for maturation of pro-neuropeptides (Kass et al., 2001). Unexpectedly, mutations in both alleles of *egl-3, n589* and *ok979*, caused weaker 30 V ON response and, moreover, much longer persistence of the speed increase after the electric shock in ON and OFF responses (Fig. 5 and 6, panels D-F), indicating that the persistent activity in the neural circuit for speed increase is down-regulated by neuropeptide signaling in wild-type animals.

## DISCUSSION

In the present study, we revealed that *C. elegans* exhibits a persistent speed increase in response to AC stimuli. This behavioral response is characterized by persistence, scalability, and valence, suggesting that it reflects an emotional state of *C. elegans*, which has never been reported. In addition, genetic analysis revealed that genes involved in electro-sensation in cartilaginous fishes and a neuropeptide biosynthesis gene are required for the response, demonstrating that the AC response of the animals is an ideal paradigm to genetically dissect the mechanism of electro-sensation as well as that of persistent behavioral states.

### Response to electric stimulus and its mechanisms in *C. elegans* and other animal species

In neuroscience research, electricity is used as an unconditioned stimulus with negative valence to cause associative learning in rodents and in flies (Quinn et al., 1974; Rescorla, 1968). In nature, however, multiple animal species are known to respond to electricity for survival purposes, such as communication, navigation and/or prey detection (Crampton, 2019; Pettigrew, 1999). For example, weakly electric African fish (*Gnathonemus petersii*) utilize their epidermal electroreceptors to receive self-produced electric signals, allowing the fish to identify, locate, and examine nearby objects (von der Emde et al., 2008). In addition, platypus (*Ornithorhynchus anatinus*) detects electric signals via their duck-like bills to locate and avoid objects when navigating in the water (Scheich et al., 1986). Blind cave salamander (*Proteus anguinus*) perceives a moving back-and-forth direct-current field and its polarity via ampullary organs to survive and navigate in their environment, which is in complete darkness as their eyes are undeveloped (Istenič & Bulog, 1984; Roth & Schlegel, 1988). In invertebrates, bumblebees (*Bombus terrestris*) sense environmental electric fields via sensory hairs to make foraging decisions (Clarke et al., 2013; Sutton et al., 2016). Such wide use of electric signals in the animal kingdom suggests physiological importance of the sensation and behavioral responses to environmental electric signals although the molecular mechanisms for electric sensation have been poorly revealed.

In this study, we established an original experimental paradigm and found that *C. elegans* responds to AC electric stimulus: The animals significantly increase their movement speed during and after the stimulus for minutes. Although the animals have also been reported to respond to DC (Gabel et al., 2007), we consider that the responses to AC and DC are substantially different for the following reasons. (1) In the DC field, the animals moved at a certain angle (~4° per 1 V/cm), which was not observed in our AC stimulus (Supplementary Fig. 3). (2) Movement speed did not change with the DC stimulus (Gabel et al., 2007).

In addition, five pairs of amphid sensory neurons were involved in the DC response (Gabel et al., 2007), while mutations in genes required for sensory signaling in the amphid sensory neurons (*tax-4*, *osm-9*, *ocr-2*, and *che-2*) did not affect the AC response significantly in our study (Fig. 5 and 6), indicating that DC and AC responses utilize different sensory mechanisms. Our result also rules out the possibility that the animals respond to increased agar temperature due to the AC stimulus, because the mutation in *tax-4*, the gene essential for temperature sensation (Komatsu et al., 1996) did not affect the response. In addition, the genes required for mechano-sensation (*mec-4, mec-10*, and *trp-4*) are not required for the AC response either.

We found that VGCC and possibly BK channel as well, the voltage-gated calcium and potassium channels for electro-sensation in the cartilaginous fishes, are involved in the AC response of *C. elegans*. The involvement of different types of voltage-gated channels in the sensation of electricity suggests that this mechanism is evolutionarily conserved. It also suggests that EGL-19 and SLO-1 may function coordinately in a subset of neurons that sense electricity. Since *egl-19* and *slo-1* are widely expressed in most of the neurons as well as in muscles (Davies et al., Lee et al., 1997; Wang et al., 2001), it would be interesting to identify the neurons where these genes function to sense the electric signals.

### Electric stimulus causes persistent behavioral response

Persistent neural activity, a sustained neural activity caused by a short term stimulus, plays critical roles in brain function, such as controlling motivation, arousal, and emotion as well as working memory and decision-making, although its detailed mechanisms have not been sufficiently elucidated (Anderson, 2016; Berridge, 2004; Curtis & Lee, 2010; Major and Tank, 2004). Persistent behavioral state is caused by persistent neural activity, suggesting that genetic analysis of persistent behavioral state can reveal molecular mechanisms(s) of persistent neural activity underlying brain functions.

We unexpectedly found that *C. elegans*’ high speed response persists after electric shock. In *C. elegans*, two other types of persistent behavioral responses related to speed change have been reported. The first is that the animal’s movement speed is elevated at high O2 concentration in *npr-1(lf*) and in the Hawaiian wild isolate CB4856, which has the same amino acid variation in *npr-1* (Cheung et al., 2005). In this behavioral response, (1) the elevated speed returns rapidly to the basal speed when the high O2 is terminated, (2) the animals still recognize and aggregate at the edge of a food lawn, and (3) a mutation in the *tax-4* CNGC homolog for sensory depolarization abolishes the response (Coates & de Bono, 2002). Another type of persistent behavioral response is roaming (Flavell et al., 2020; Fujiwara et al., 2002). Roaming is a behavioral state with high movement speed, although it is only exhibited when the animals are on food and requires serotonin signaling. Because the behavioral response to electric shock persists more than 2 min after 30-45 sec stimulus with 75 V and more than 1.5 min after only 5 sec stimulus, is not affected by food stimulus, and does not require CNGC activity or serotonin signaling, electric shock response is likely different from the above-mentioned two behavioral responses, and its analysis may provide a unique opportunity for genetic dissection of a persistent behavioral state and neural activity.

The 30 or 75 V of voltage used in this study may appear artificial. However, we consider that the responses of *C. elegans* to these stimuli reflect physiologically meaningful biological mechanisms for the following reasons: (1) The range of voltage per length (30–75 V/6 cm = 5–12.5 V/cm) is similar to the one used to study the animal’s DC response (3–12 V/cm) (Gabel et al., 2007). (2) The electric current flowing inside the worm’s body could be weak because it depends on the resistance of its body and cuticle. (3) Only a 5 second stimulus causes a persistent response that lasts more than a minute, meaning that the electric shock itself is just a trigger and what we observe is a physiological response to that trigger. (4) Fear conditioning in rodents is also triggered by electrical shock. The speed increase behavior we observed may resemble fleeing, one of the most common responses caused by fear in higher animals and humans (Adolphs, 2013; Bliss-Moreau, 2017; Mobbs & Kim, 2015).

### Response to the electric stimulus may reflect a form of emotion

Emotions are internal brain states triggered by certain types of environmental stimuli, which are associated with cognitive, behavioral, and physiological responses (Abbott, 2020; Anderson & Adolphs, 2014; Nettle & Bateson, 2012; Perry & Baciadonna, 2017). Recently, multiple species of invertebrates are considered to possess internal brain states that resemble what we consider to be emotions (Bacqué-Cazenave et al., 2017; Cwyn et al., 2016; Gibson et al., 2015; Hamilton et al., 2016; Mohammad et al., 2016; Pascal et al., 2014). One of the most prominent characteristics of emotion across animal species is its persistence: For example, even a transient environmental stimulus can cause a persistent behavioral response, such as courtship, aggressive, and defensive behavior (Abbott, 2020; Anderson & Adolphs, 2014; Nettle & Bateson, 2012; Paul & Mendl, 2018; Perry & Baciadonna, 2017).

Anderson and Adolphs proposed a new framework to study emotions across animal species, wherein hallmarks of an emotional state are persistence, scalability, valence, and generalization. In addition to persistence (Fig. 4A–C), we also consider that the electric response has negative valence. This is because the animals ignore food during the electric shock response (Fig. 4D and E, and Sup. Fig. 5), despite the fact that food is one of the most influential signals for *C. elegans*, affecting many aspects of their behavior. For example, during the high speed state caused by high O2, animals still recognize and stay at the edge of a food lawn (Cheung et al., 2005; Coates & de Bono, 2002), suggesting that the electric shock signal has a strong negative valence that overrides the strong positive valence of food. The third point is the scalability—stronger stimulus causes stronger behavioral response. Compared to the 30 V stimulus, the 75 V stimulus results in a larger number of immobile animals during the stimulus period (right panels in Fig. 2A and B) as well as a longer-lasting high speed response after the stimulus (Fig. 4A and C). The fourth point is generalization – the same emotional state can be triggered by different stimuli and, in turn, the emotional state triggered by one stimulus can then affect responses to other stimuli. The lack of response to food during and following our electric stimulus might supports this point as well, as the emotional state induced by electricity influences the response to food, an entirely different stimulus.

Taken together, these results suggest that the animal’s response to electric shock represents a form of emotion, probably fear. As we revealed that the persistent aspect of the behavioral response is regulated by neuropeptide signaling (Fig. 5 and 6), which may resemble the neuropeptide regulation of fear in mammals including humans (Bowers et al., 2012; Comeras et al., 2019; van den Burg & Stoop, 2019), the fear-like brain state may be regulated by evolutionarily conserved molecular mechanisms.

In summary, we found that *C. elegans* persistently responds to electric shock, which is regulated by voltage-gated ion channels and neuropeptide signaling. Our findings may suggest the following model (Fig. 7). When the animals sense 30 or 75V AC stimulus at 4 Hz, the stimulus is sensed with the VGCC and BK channel and their internal state transits from basal speed state to persistent high speed state. The persistent high speed state eventually returns to the basal speed state, which requires neuropeptide signaling. By taking advantage of connectome information and the methods for imaging whole brain activity of identified neurons (Randi & Leifer, 2020; Wen et al., 2021; White et al., 1986; Yemini et al., 2021), *C. elegans* may become one of the ideal models for revealing the dynamic information processing involved in the entire neural circuit that regulates emotion.

**Fig. 7.**
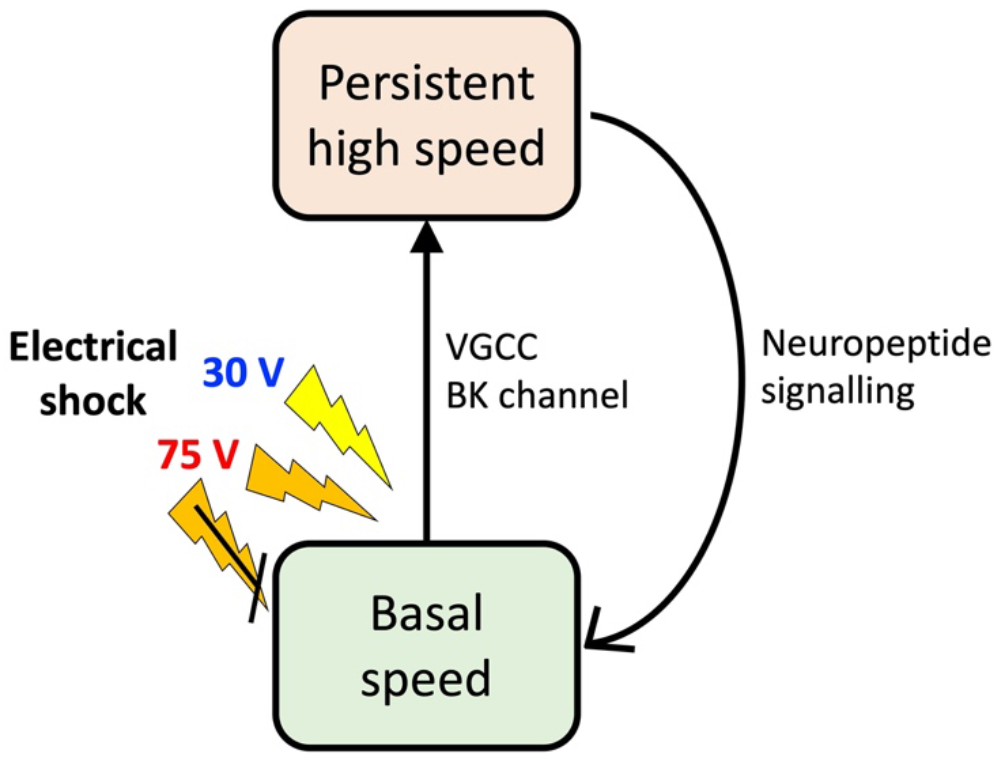
Model for mechanism of speed increase caused by electric shock.

## MATERIALS AND METHODS

### *C. elegans* strains

*C. elegans* strains were maintained with standard procedures (Brenner, 1974). In brief, for regular cultivation, animals were grown on standard 6 cm nematode growth medium (NGM) agar plates which had been spread with *E. coli* strain OP50 and incubated at 19.0-19.5 °C. Strains used were the wild-type strain Bristol N2 and mutant strains PR678 *tax-4(p678)*, CX4652 *osm-9(ky10)*;*ocr-2(ak47)*, CB1033 *che-2(e1033)*, TU253 *mec-4(u253)*, ZB2551 *mec-10(tm1552)*, TQ296 *trp-4(sy695)*, MT1212 *egl-19(n582)*, DA995 *egl-19(ad995)*, JD21 *cca-1(ad1650)*, CB55 *unc-2(e55)*, VC854 *unc-2(gk366)*, NM1968 *slo-1(js379)*, BZ142 *slo-1(eg142)*, KDK11 *cat-2(tm2261)*, MT7988 *bas-1(ad446)*, GR1321 *tph-1(mg280)*, RB993 *tdc-1(ok914)*, VC671 *egl-3(ok979)*, and MT1219 *egl-3(n589)*.

### *C. elegans* cultivation for electric shock behavioral assay

Before the behavioral assay, animals were cultivated as described previously (Kimura et al., 2010). In brief, four adult wild-type animals were placed onto NGM agar plates with OP50 and kept at 19.5°C for 7.5 hours before being removed. After removal, these plates were incubated at 19.0–19.5 °C for 3 days until the assay day. On the assay day, about 100 synchronized young adult animals were grown on each plate. As some mutant animals had slower growth or laid fewer eggs than wild-type animals did, the incubation temperature and number of these mutant animals were adjusted and increased accordingly in order to obtain a comparable developmental stage (i.e. young adult) and worm number with the wild-type animals. All behavioral assays were carried out with young adult hermaphrodites.

### Experimental instruments for electric shock behavioral assay

The following electric instruments (Fig. 1) were utilized for the electric shock behavioral assay. A 50 MHz Arbitrary Waveform Generator (FGX-295, Texio Technology Corporation) was used to generate different types of electric waveforms over a wide range of frequencies. This waveform generator has an output limit of 10 V. Thus, an AC Power Supply (PCR500MA, Kikusui Electronics Corp.) was used to amplify the voltage supply. We also used a Digital Storage Oscilloscope (DCS-1054B, Texio Technology Corporation) in parallel to measure the voltage and observe the electric waveforms produced as well as a Digital Multimeter (PC720M, Sanwa Electric Instrument Co., Ltd.) to measure current. A USB camera (DMK72AUC02, The Imaging Source Co., Ltd.) with a lens (LM16JC5M2, Kowa) was used to record trajectories produced by the animals.

### Electric shock behavioral assay with small OP50 bacterial food patch

Most of the behavioral assays were conducted on 9 cm NGM agar plates seeded with a small food patch unless indicated otherwise. For the food patch, the bacteria OP50 was grown in 100 mL of LB culture overnight at 37°C, spun down and resuspended in 10 volumes of NGM buffer, and 5 μL of the suspension was applied at the center of the plate to create a food patch 3 × 10 mm in size on the assay day. This process was used to minimize the thickness of the food patch as it prevents clear images of worms in the patch. Four animals per plate were placed in the food patch one hour before the assay to accustom the animals to the environment and to reduce their movement speed to the basal level. The assay plates were then inverted and placed onto a custom-made copper plate bridge, whose distance is 6 cm (Fig. 1). The images were acquired 2 frames per s, and electric shock was delivered with the conditions described in each figure. Move-tr/2D software (Library Inc., Japan) was used to calculate the *x-y* coordinates of the animal centroids in each image frame, which were then analyzed using Excel (Microsoft) or R (The R Project) to calculate the animal’s speed. In general, the moving median for ±1 frame was calculated to remove noise for each animal and then ensemble averaged for each condition. Baseline speed was calculated from the average speed over 30 s before the stimulation, and ΔSpeed was calculated by subtracting the baseline value from each animal’s speed during or after the stimulus.

### Electric shock behavioral assay with full or strip-like OP50 bacterial food lawn

For the assays conducted with full food lawn, the area of assay plates between the copper plates were fully seeded or seeded in a stripe-shape with OP50 and kept on the bench overnight until the assay began. Animals grown in regular cultivation plates were washed in two droplets of NGM buffer and then transferred to the center of the assay plate and left for 5 minutes. The rest of the procedures were the same as for assays conducted with small food patch.

To detect the outward and inward movement on the food stripes (Supplementary Fig. 5), the food positions were first indicated on each image series by the experimenter and the animal’s centroid across the boundary was automatically calculated by a custom-made program.

### Investigation of relationship among speed increase, current and voltage

Three different types of NGM agar plates were prepared with varying salt concentrations and similar osmolarities: High-salt plates had 200 mM sodium chloride; low-salt plates had 10 mM sodium chloride and 380 mM sucrose; control plates had 50 mM sodium chloride and 300 mM sucrose. The purpose of adding sucrose into the plates was to adjust and balance the osmolarity. The final total osmolarity for sodium chloride (Na^+^ and Cl^-^) and sucrose for all the plates was 400 mOsm. The rest of the procedures were the same as for assays conducted with small food patch.

### Data analysis and statistics

All the statistical analyses were performed in R (The R Project). Generally, data of 20 – 50 animals in total from 9 plates from 3 days of experiments for each condition were pooled and analyzed together. We chose this sample number based on a large scale behavioral analysis of *C. elegans* (Yemini et al., 2013). Data is presented as means ± SD unless otherwise specified. Experimental conditions, such as the electric stimulation or different strains were randomized on a daily basis.

## ACKNOWLEDGEMENT

We thank Liting Chen for having provided the idea of “worm’s emotion” for K.D.K., Yuki Tanimoto and Yuka Tsuda for the initial phase of the electric shock paradigm, Shinobu Aoyagi for setting up the system, Chentao Wen for developing the outward/inward scoring program, and Young-Jai You, Aki Takahashi, Atsuko Nakagawa and the Kimura lab members for their valuable advice, comments and technical assistance for the study. Nematode strains were provided by the Caenorhabditis Genetics Center (funded by the NIH Office of Research Infrastructure Programs P40 OD010440).

## FUNDING

This study was supported by Japan Society for the Promotion of Science (KAKENHI JP16H06545, 20H05700, 21H00448 and 21K19274 to K.D.K.), Grant-in-Aid for Research in Nagoya City University (48, 1912011, 1921102 and 2121101), the Joint Research by National Institutes of Natural Sciences (01112002), Toyoaki Scholarship Foundation, and RIKEN Center for Advanced Intelligence Project (to K.D.K). L.F.T. was supported by Japanese Government (MEXT) Scholarship.

## COMPETING INTERESTS

The authors declare no competing interests.

## Notes

### Competing Interest Statement

The authors have declared no competing interest.

