## Supplementary Figures for "Electric shock causes a fear-like persistent behavioral response in the nematode *Caenorhabditis elegans*"

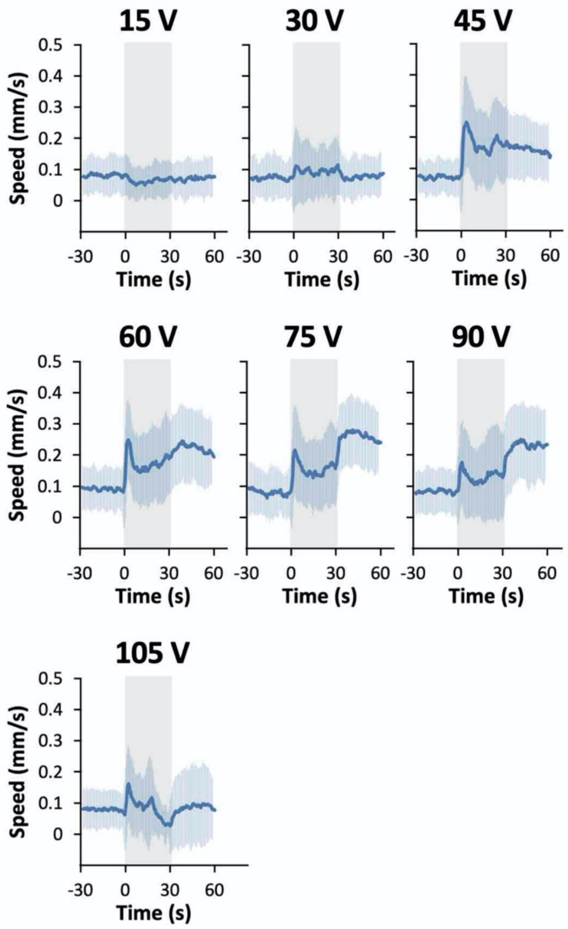


**Supplementary Fig. 1**. Speed-time graphs with different voltage stimulation at 60 Hz. Gray indicates the duration of electric stimulation (0-30 s). The thick line and the shaded region indicate the average ± SD. Sample numbers were 57–58 per condition, and the details are described in the Supplementary Table.

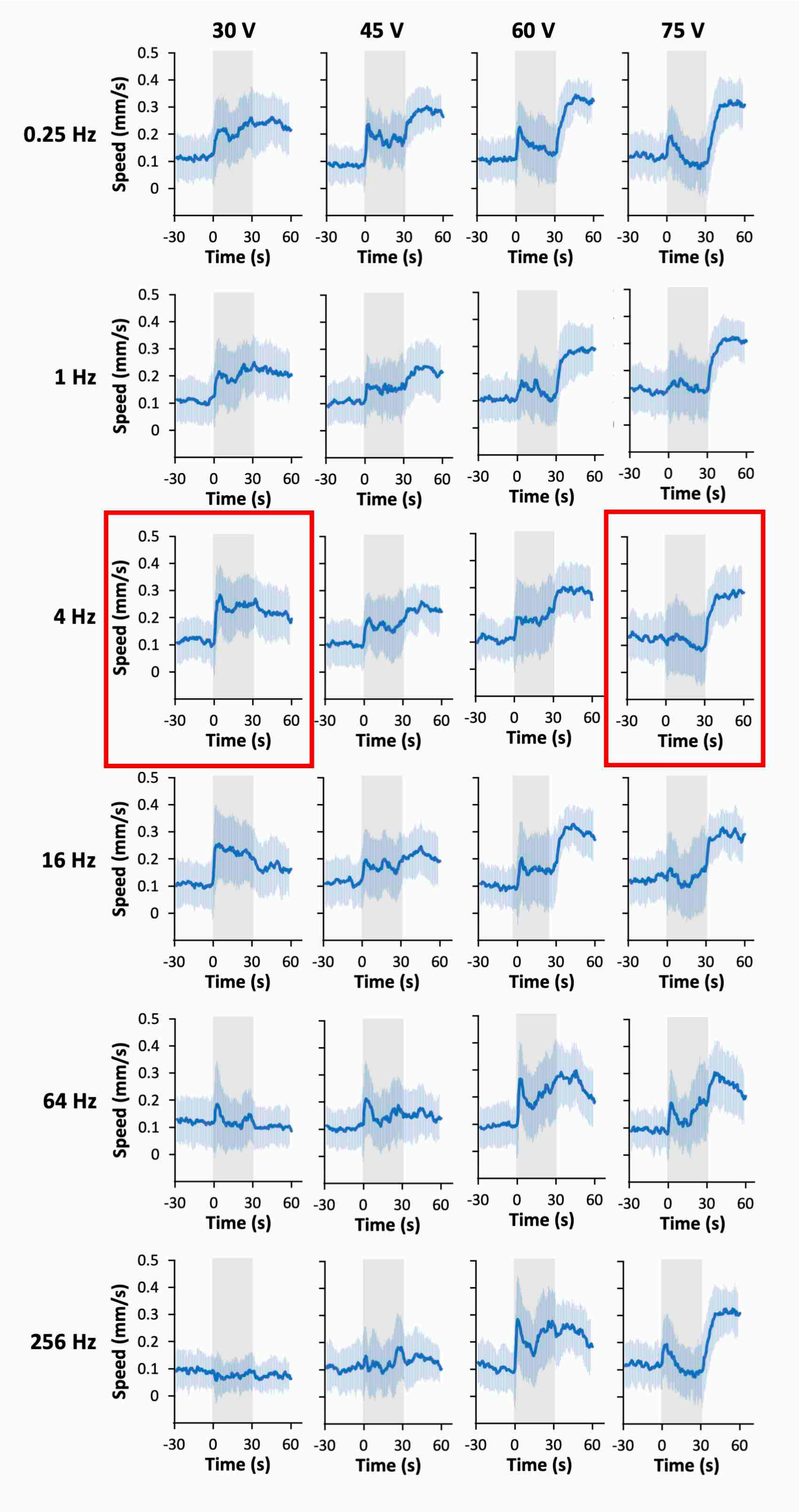


**Supplementary Fig. 2**. Speed-time graphs with different voltage stimulation at different frequencies. Gray indicates the duration of electric stimulation (0-30 s). The thick line and the shaded region indicate the average ± SD. 30 and 75 V at 4 Hz (red rectangles) were chosen for further analyses. Sample numbers were 33–37 per condition, and the details are described in the Supplementary Table.

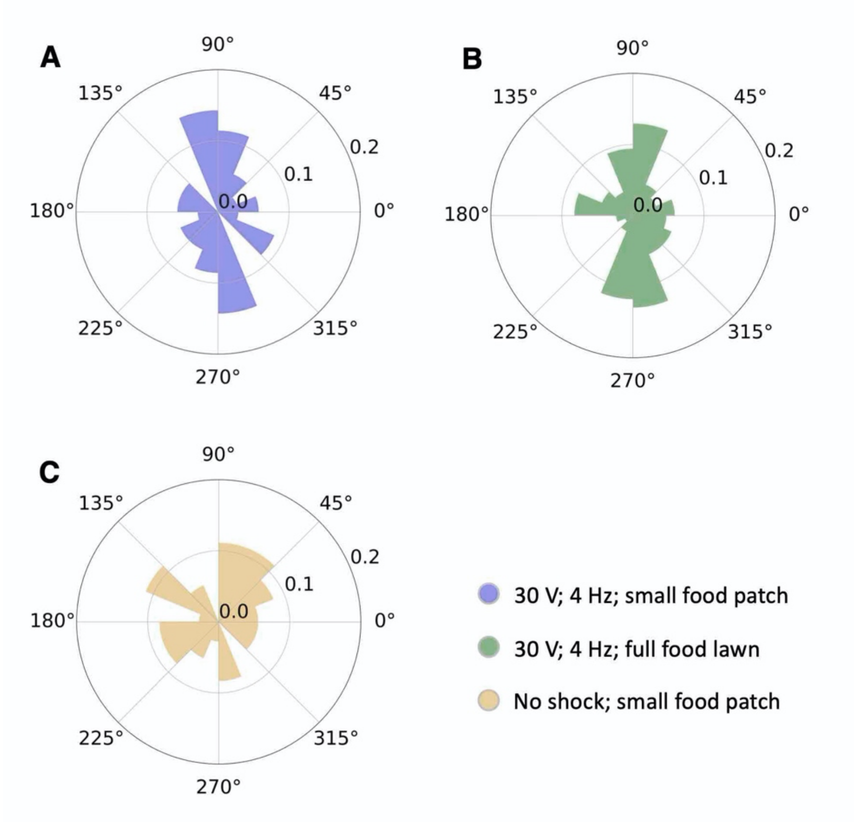


**Supplementary Fig. 3**. Movement directions of animals during the response. The angles of movement vectors from the beginning to the first 2 min of the stimulation were plotted. **A-C,** Rose plot for animals which were assayed on plate with small food patch (**A**, n = 35; Group 1) or full food lawn (**B**, n = 85; Group 2) with 30 V at 4 Hz, or small food patch without electric stimulation (**C**, n = 36; Group 3). Bin number for each chart is set at 16 bins. Statistical analysis performed is Watson U2 test, and *p* values for Groups 1 vs 2, 1 vs 3, and 2 vs 3 were all >0.05.

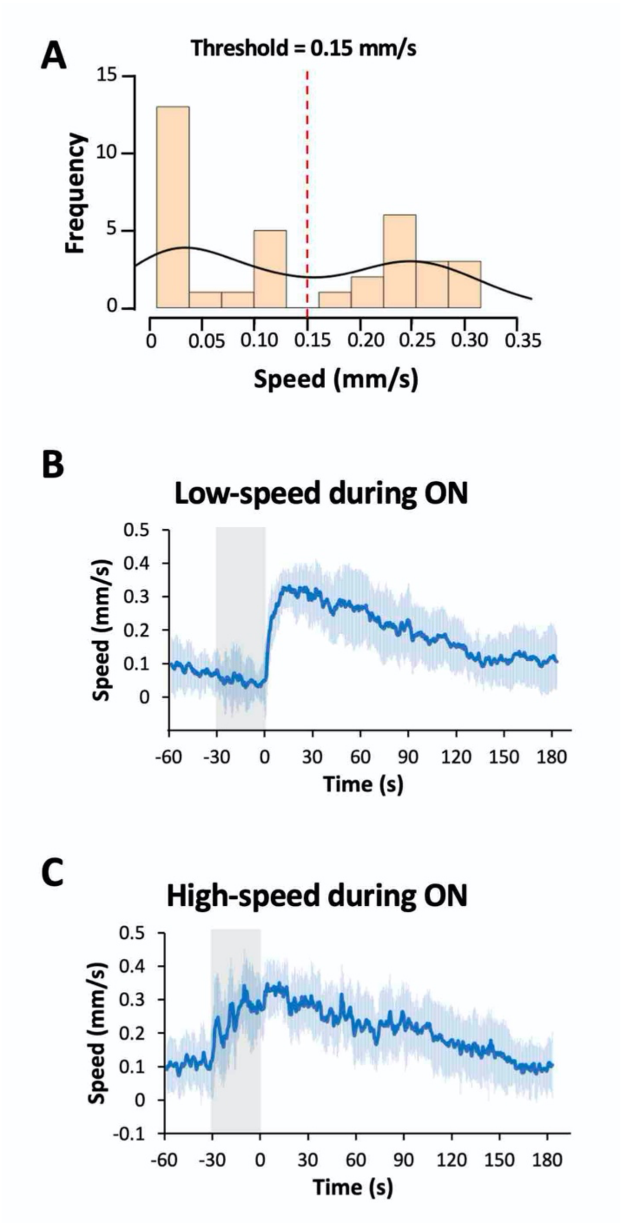


**Supplementary Fig. 4**. Low and high speed groups during 75 V stimulation. **A**, Histogram and its density (black line) indicates speed of each animal during the electric shock. From the histogram, we set the threshold at 0.15 mm/s to separate the low- (**B**) and high-speed (**C**) groups. Sample numbers were 20 and 15 for lower and higher speed groups, respectively, and the details are described in Supplementary Table.

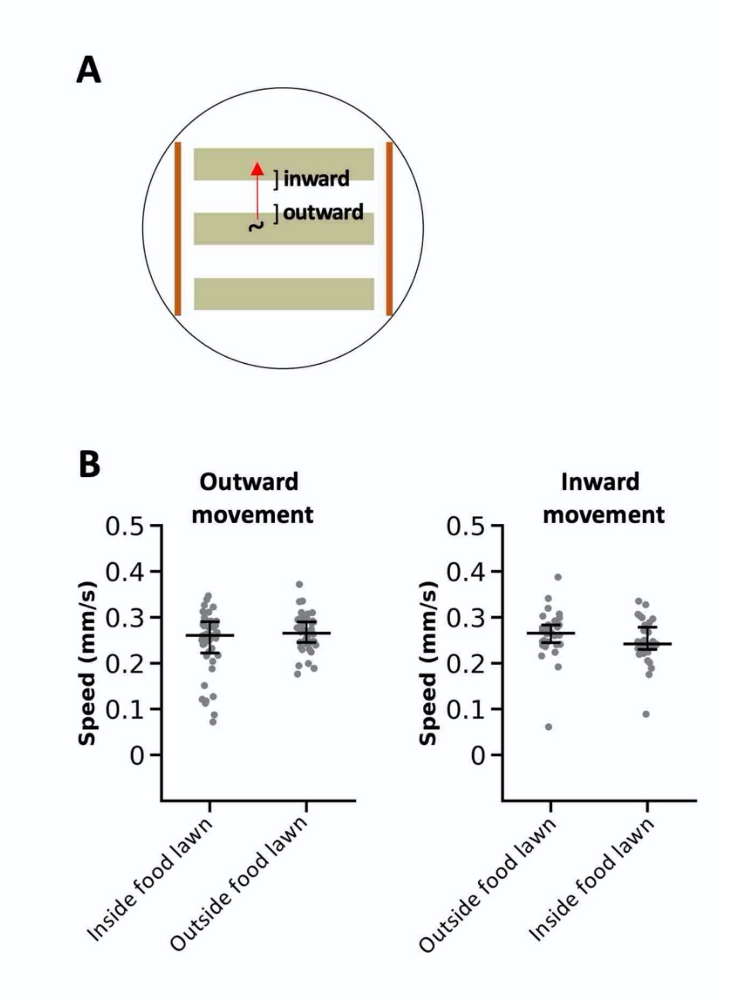


**Supplementary Fig. 5.** Worms' speed did not change when they move in or out of food. **A,** Illustration showing worms' movement across multiple food strips. When worms leave food strip and enter no food area, this movement is defined as "outward movement". When worms enter food strip from no food area, this movement is defined as "inward movement". **B,** Scatter plot showing average speed of individual animals with outward (left; n = 44) or inward (right; n = 32) movement during 30 V stimulation for 4 min. The average speed was calculated 10 s before and after the food exit/entry. Statistical analysis was performed by Wilcoxon signed-rank test, and no significant difference was observed.

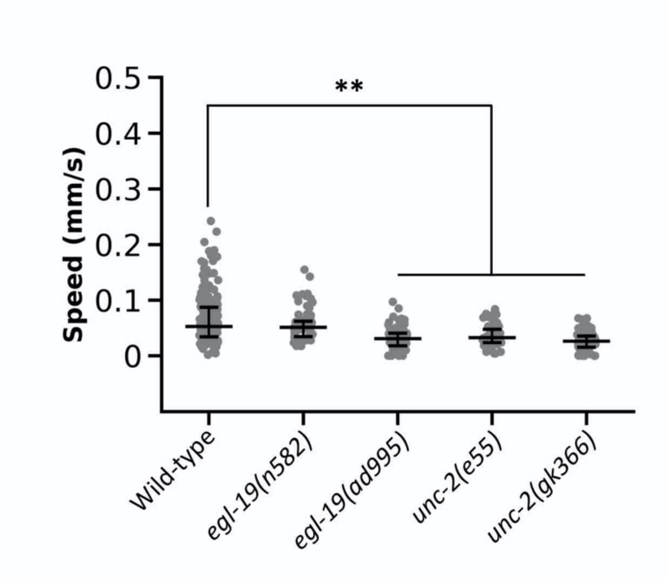


**Supplementary Fig. 6.** Basal speeds of wild-type and VGCC mutants before the 30 V and 75 V stimulations. Statistical values were calculated using Kruskal-Wallis test with Bonferroni correction. ** $p < 0.001$.

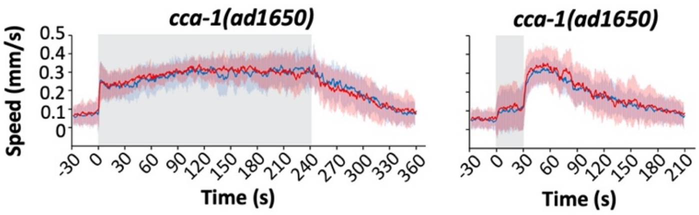


**Supplementary Fig. 7.** Speed-time graphs of ON (30 V, left) and OFF (75 V, right) responses of *cca-1*.

**Supplementary Video 1.** Responses of animals in one individual assay plate subjected to 30 V stimulation at 4 Hz for 30 seconds. The trajectories and corresponding individual speed values produced by the animals were shown. The custom-made copper plate bridges were placed at left and right side of field view. Video speed is adjusted to 2X.

**Supplementary Video 2.** Responses of animals in one individual assay plate subjected to 30 V stimulation at 4 Hz for 30 seconds. This video is from raw images used for the video 1. Video speed is adjusted to 2X.

**Supplementary Video 3.** Responses of animals in one individual assay plate subjected to 75 V stimulation at 4 Hz for 30 seconds. The trajectories and corresponding individual speed values produced by the animals were shown. The movement of all animals in this plate are completely suppressed during 75 V stimulation, but this phenomenon is not applicable in every plate as some animals are still able to move during this stimulation period. Similarly to the video 1, the custom-made copper plate bridges were placed at left and right side of field view. Video speed is adjusted to 2X.

**Supplementary Video 4.** Responses of animals in one individual assay plate subjected to 75 V stimulation at 4 Hz for 30 seconds. This video is from raw images used for the video 3. Video speed is adjusted to 2X.
